# Olfaction regulates peripheral mitophagy and mitochondrial function

**DOI:** 10.1101/2023.08.21.554156

**Authors:** Julian G. Dishart, Corinne Pender, Koning Shen, Hanlin Zhang, Megan Ly, Madison Webb, Andrew Dillin

## Abstract

The central nervous system is a master regulator of peripheral homeostasis and cellular-stress responses; however, the contexts for which this regulatory capability evolved remain unknown. The olfactory sensory nervous system has access to privileged information about environmental conditions and can signal to the periphery to prepare for potential metabolic perturbations. The unfolded protein response of the mitochondria (UPR^MT^) is upregulated upon infection by many pathogens and in metabolic flux, and pathogenic infection and metabolic byproducts are a present hazard in consuming nutrients. Therefore, we asked whether the olfactory nervous system in *C. elegans* regulates the UPR^MT^ cell nonautonomously. We found that loss of a single olfactory neuron pair, AWC, led to robust induction of the UPR^MT^ downstream of enhanced, serotonin-dependent mitophagy. Further, AWC ablation confers resistance to the pathogenic bacteria *Pseudomonas aeruginosa* partially dependent on the UPR^MT^ transcription factor *atfs-1*, and fully dependent on mitophagy machinery *pdr-1/Parkin*. These data demonstrate a novel role for the olfactory nervous system in regulating whole-organism mitochondrial dynamics, perhaps in preparation for postprandial metabolic stress or pathogenic infection.

## INTRODUCTION

Coordinating stress responses across tissues is paramount for maintaining cellular homeostasis and longevity in response to environmental insults. A robust body of literature has established the central nervous system as a master regulator of cellular stress responses across the whole organism. For example, activating the unfolded protein response of the endoplasmic reticulum (UPR^ER^) (*1*), the heat shock response (*2*), and the mitochondrial unfolded protein response (UPR^MT^) (*3*, *4*) in neurons results in cell nonautonomous induction of their respective stress response in peripheral tissues. This nonautonomous coordination promotes survival in aging and in conditions of cellular stress, leading to the hypothesis that the nervous system evolved the ability to prepare the organism to withstand noxious environmental conditions. More recently, work on the nonautonomous regulation of UPR^MT^ has focused on the neural circuits required to drive the UPR^MT^ and other mitochondria-associated phenotypes under conditions of neuronal mitochondria dysfunction. For example, induction of mitochondrial stress in neurons by knockout of FZO-1/mitofusin drives peripheral UPR^MT^ induction, enhances mitophagy, and increases pathogen resistance dependent on serotoninergic, tyraminergic, and neuropeptidergic signaling (*5*). Despite the insights into the signaling cascades and neuronal players required for peripheral induction of the UPR^MT^ by the central nervous system, the evolutionary context and sensory modalities that necessitate this coordination remains incompletely understood.

The sensory nervous system, particularly the olfactory nervous system, has emerged as a nonautonomous regulator of metabolic homeostasis and lifespan. In mice, olfactory exposure to food activates hypothalamic neurons, signaling to the liver to activate the ER^UPR^ and to initiate lipid synthesis (*6*), while olfactory neuron ablation abates diet-induced obesity and increases mitochondrial respiration in fat tissue through noradrenergic afferent signaling (*7*). A connection between olfaction and metabolic regulation has also been established in the model organism *C. elegans*, suggesting an evolutionarily conserved role for olfactory circuits in regulating peripheral physiology. Loss of function in distinct olfactory neurons extends lifespan by downregulating insulin signaling (*8*), and silencing a single olfactory neuron pair, AWC, regulates lipid metabolism and enhanced proteostasis through neuroendocrine control (*9*, *10*). These studies indicate that the olfactory nervous system plays a role in assessing environmental nutritional status, and signals to the periphery to prepare for food intake.

All organisms must consume nutrients to survive; however, eating poses significant cellular hazards. In eating, tissues are not only tasked with metabolizing nutrients, but must also withstand and clear toxic metabolites, like mitochondria-generated reactive oxygen species (mtROS) and must defend against food-borne pathogens. Pathogenic infection is a formidable cellular stressor, and organisms, including the bacterivore *C. elegans,* induce various genetic programs to counteract pathogen-associated damage. For example, during infection with the pathogenic bacteria *Pseudomonas aeruginosa* (*P. aeruginosa*), *C. elegans* activate the UPR^MT^ transcription factor *atfs-1* to resolve mitochondrial damage and to induce innate immune response genes (*11*). Loss of *atfs-1* activity and diminished UPR^MT^ induction results in sensitivity to *P. aeruginosa* infection (*11*). In their natural milieu, *C. elegans* are exposed to myriad commensal and pathogenic bacteria and must discern which are helpful or harmful, primarily through the detection of bacterial metabolites. Olfaction is a primary mode of detection of such volatile metabolites, meaning that olfactory neurons have privileged access to information about potential pathogenic threats in the environment. We hypothesize that olfactory neurons might be top-most coordinators of cellular stress-response induction and may serve as an early warning system to prepare the periphery for pathogenic infection. Indeed, recent work in *C. elegans* has shown that loss of certain sensory neurons, particularly the olfactory neuron AWC, contributes to *P. aeruginosa* resistance; however, whether olfactory regulation of peripheral stress responses is responsible for this resistance has been left unexplored (*12*).

AWC neurons are tonically active in the absence of odorants and become silenced when odorants are presented. AWC neurons inhibit their downstream interneuron partners; therefore, when ablated, AWC-regulated circuitry is de-repressed (*13*). Here, we show that ablation of the olfactory neuron pair AWC is sufficient to drive peripheral activation of UPR^MT^, reduce organism-wide mitochondrial oxidative phosphorylation (OXPHOS) rates, and reduce organismal mitochondrial DNA (mtDNA). These phenotypes are dependent on serotonergic signaling, suggesting that AWC-mediated olfactory circuitry regulates peripheral UPR^MT^ and mitochondrial mass cell nonautonomously. Moreover, induction of the UPR^MT^, reduction in OXPHOS, and reduction in UPR^MT^ downstream of AWC ablation depends on the mitophagy machinery *pdr-1* (PRKN homolog). Finally, we find that AWC-mediated *P. aeruginosa* infection resistance is partially dependent on the UPR^MT^ transcription factor *atfs-1* and fully dependent on *pdr-1,* indicating that olfactory neurons can prepare the organism for pathogenic insult through regulation of mitochondrial dynamics.

## RESULTS

### Ablation of olfactory neuron AWC induces the *UPR^MT^*, reduces mitochondrial oxidative phosphorylation, and reduces mitochondrial mass

*C. elegans* employ three primary olfactory neuron pairs to coordinate behavioral responses to volatile odorants. Two of these neuron pairs, AWA and AWC, mediate chemotaxis toward appetitive olfactory cues, while AWB neurons initiate chemotaxis away from noxious cues (*14*). AWC neurons are unique in that they are constitutively active in the absence of food odors, and tonically exert inhibitory control over their downstream interneuron partners. On the other hand, in the presence of its odorant ligands, AWC neurons are silenced and release this inhibitory regulation (*13*). Therefore, ablation of AWC neurons mimics odor presentation, leading to chronic disinhibition of AWC-regulated neural circuitry.

To understand whether olfactory neurons regulate peripheral mitochondrial dynamics, we exploited the neurobiology of AWC neurons, and first investigated whether *hsp-6* expression, a mitochondrial chaperone protein and marker for induction of the mitochondrial unfolded protein response (UPR^MT^) is induced in AWC-ablated [AWC(-)] animals (**Fig. 1A**). To ablate AWC, we used a previously validated transgenic strain that expresses cleaved caspase under the AWC-specific promoter, *ceh-36* (*ceh-36p::caspase*) (*15*) and crossed them to the UPR^MT^ transcriptional reporter *hsp-6::gfp*. We found that AWC(-); *hsp-6::gfp* showed pronounced activation of the UPR^MT^ (**Fig. 1 B and C**).

**Figure 1:**
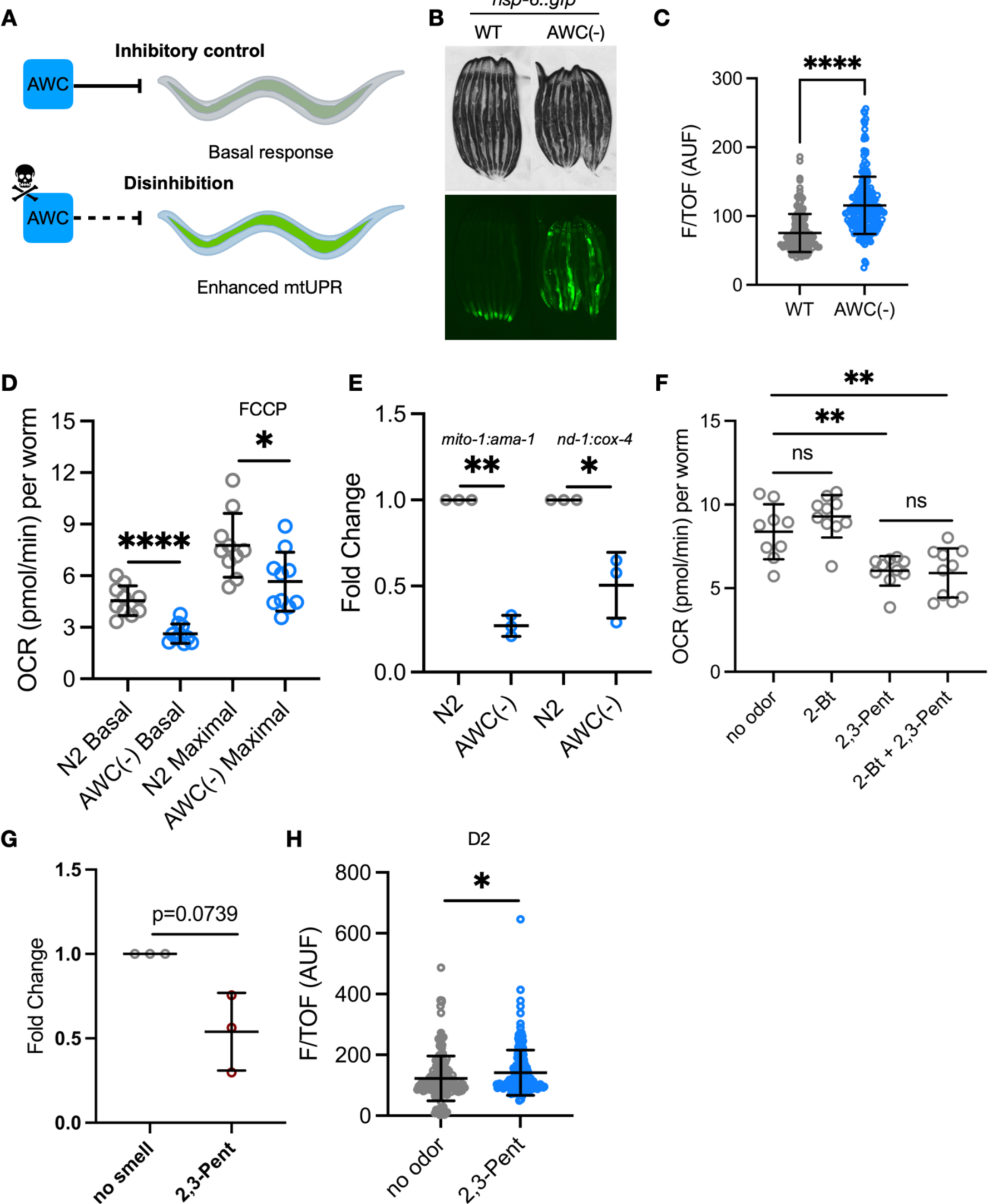
AWC ablation induces the UPR^MT^, reduces organismal OXPHOS, and reduces mtDNA. (A) Schematic illustrating hypothesis that ablation of AWC disinhibits downstream circuitry to induce peripheral UPR^MT^. (B) Representative fluorescent images of *hsp-6::gfp* in N2 and AWC(-) animals. (C) Integrated fluorescence intensity measured by bioSorter of hsp-6::GFP of strains imaged in panel A. AUF, arbitrary units of fluorescence. Two-tailed unpaired t test with Welch’s correction, ****P<0.0001. (D) Oxygen consumption rate (OCR) without (basal) and with FCCP (maximal). Two-tailed unpaired t test with Welch’s correction, ****P<0.0001, *P<0.05. (E) Log2 fold change of the ratio of mtDNA to genomic DNA in AWC(-) normalized to N2, measured by qPCR. N = 3 biological replicates.Two-tailed unpaired t test with Welch’s correction, **P<0.01, *P<0.05. (F) OCR in N2 worms treated with 2-Bt, 2,3-Pent, and 2-Bt with 2,3-Pent. Two-tailed unpaired t test with Welch’s correction, P=0.1936, **P<0.001, P=0.8133. (G) Log2 fold change of the ratio of mtDNA (*mito-1*) to genomic DNA (*ama-1*) in 2,3-Pent treated WT animals normalized to no odor treated WT animals, measured by qPCR. N = 3 biological replicates. Two-tailed unpaired t test with Welch’s correction, P=0.0739. (H) Integrated fluorescence intensity measured by bioSorter of hsp-6::GFP in no odor and 2,3-Pent treated WT animals. AUF, arbitrary units of fluorescence. Two-tailed unpaired t test with Welch’s correction, *P<0.05.

Curious whether the induction of UPR^MT^ downstream of AWC circuit disinhibition correlated with changes in mitochondrial function, we measured oxygen consumption rates (OCR), a readout of mitochondrial oxidative phosphorylation (OXPHOS), in wild-type N2 animals versus AWC(-) animals at the last larval stage (L4). We assayed mitochondrial phenotypes at the L4 stage to assess somatic changes in mitochondrial function, before the expansion of the germline, which contributes to the bulk of *C. elegans* mitochondrial content (*16*). We found that AWC(-) animals had markedly reduced OCR at baseline, and also when treated with the OXPHOS uncoupler FCCP (maximal OCR) (**Fig. 1D)**. The proportional reduction in basal and maximal OCR to suggest that AWC(-) animals have lower mitochondrial mass. To test this, we compared the ratio of mitochondrial genomic genes to nuclear genomic housekeeping genes (*17*). We found that the ratio of *mito-1* to *ama-1* and the ratio of *nd-1* to *cox-4* was significantly reduced in AWC(-) animals compared to N2 at the L4 stage, indicating a significant reduction in mitochondrial DNA (mtDNA) (**Fig. 1E**). To assess whether these effects were dependent on the germline, we assayed OCR and *hsp-6* expression in day 1 (D1) AWC(-) and N2 adults treated with or without FUDR. FUDR is a chemical inhibitor of DNA synthesis that blocks germline production in post-mitotic *C. elegans*. We found that UPR^MT^ expression increased and OCR decreased in D1 AWC(-) mutants with and without FUDR, suggesting that these phenotypes do not depend on the germline and persist into adulthood (**Fig. S1, A and C**).

Next, we were interested in whether genetic or naturalistic silencing of AWC recapitulates the changes in mitochondrial dynamics achieved through ablation of AWC. First, we utilized a transgenic mutant that expresses a histamine-gated chloride channel under the AWC-specific promoter *ceh-36* (*ceh-36p::HisCl*) to probe the mitochondrial phenotypes (*18*). With this strain, histamine (HA) treatment silences AWC through hyperpolarization, while HA alone has no known endogenous function in *C. elegans* (*18*). We found that *ceh-36p::HisCl; hsp-6::gfp* animals had increased UPR^MT^ expression when treated with HA (**Fig. S1, D and E**), and that HA-treated *ceh-36p::HisCl* animals had reduced OCR and mtDNA (**Fig. S1, F and G**). Together, these data indicate that disinhibition of AWC-mediated circuitry upregulates the UPR^MT^, reduces OXPHOS, and reduces mtDNA.

AWC neurons develop asymmetrically into two subtypes (AWC^ON^ and AWC^OFF^), which diverge through differential transcriptional control (*19*). These subtypes express shared and distinct odorant receptors, whereby AWC^ON^ is silenced by the volatile bacterial metabolite butanone (Bt) and AWC^OFF^ is silenced by the volatile bacterial metabolite 2,3-pentanedione (2,3-Pent) (*19*). We exposed N2 wild-type animals to Bt, 2,3-Pent, and a combination of 2-Bt and 2,3-Pent, and found that 2,3-Pent exposed animals showed reduced OXPHOS (**Fig. 1F)**. Notably, there was no combinatorial effect of 2,3-Pent and Bt on OCR (**Fig. 1F**). 2,3-Pent exposure failed to significantly decrease mtDNA compared to untreated N2 animals, but exhibited a trending reduction in mtDNA (**Fig. 1G**). Interestingly, 2,3-Pent was insufficient to drive induction of UPR^MT^ at D1 (data not shown), but mildly induced *hsp-6* expression from chronic exposure through day 2 (D2) of adulthood (**Fig. 1H**). We interpreted these results to suggest that silencing AWC^OFF^ by 2,3-Pent is sufficient to reduce OXPHOS, and that this change in mitochondrial function may be due to lower levels in mtDNA. Further, these phenotypes may precede activation of the UPR^MT^, as UPR^MT^ activation occurred temporally after reductions in OCR. Alternatively, silencing of AWC^OFF^ by 2,3-Pent may not be robust enough to phenocopy UPR^MT^ activation and mtDNA depletion achieved by cellular ablation.

### Ablation of AWC olfactory neurons remodels mitochondria through neurotransmission

To test whether ablation of AWC remodels mitochondrial function cell-nonautonomously, we tested the dependence of the mitochondrial phenotypes on machinery required for neurotransmission. In AWC(-); *hsp-6::gfp* animals, we examined the effect of mutation in *unc-13,* which is required for small clear vesicle (SCV) neurotransmission, and *unc-31,* which is required for dense core vesicle (DCV) neurotransmission. We found that loss of either *unc-13* or *unc-31* suppressed peripheral induction of UPR^MT^ in AWC(-); *hsp-6::gfp* reporters (**Fig. 2, A and B**). Next, we assayed OCR and measured mtDNA in AWC(-); *unc-31(e928)* compared to *unc-31(e938)* mutants, and in AWC(-); *unc-13(s69)* compared to *unc-13(s69)* mutants. We found that the reduction in both OCR and mtDNA in AWC(-) depends on either *unc-31* or *unc-13* (**Fig. 2, C - F**). These data indicate that ablation of AWC drives SCV and DCV signaling to the periphery to induce the UPR^MT^, reduce OCR, and reduce mtDNA.

**Figure 2.**
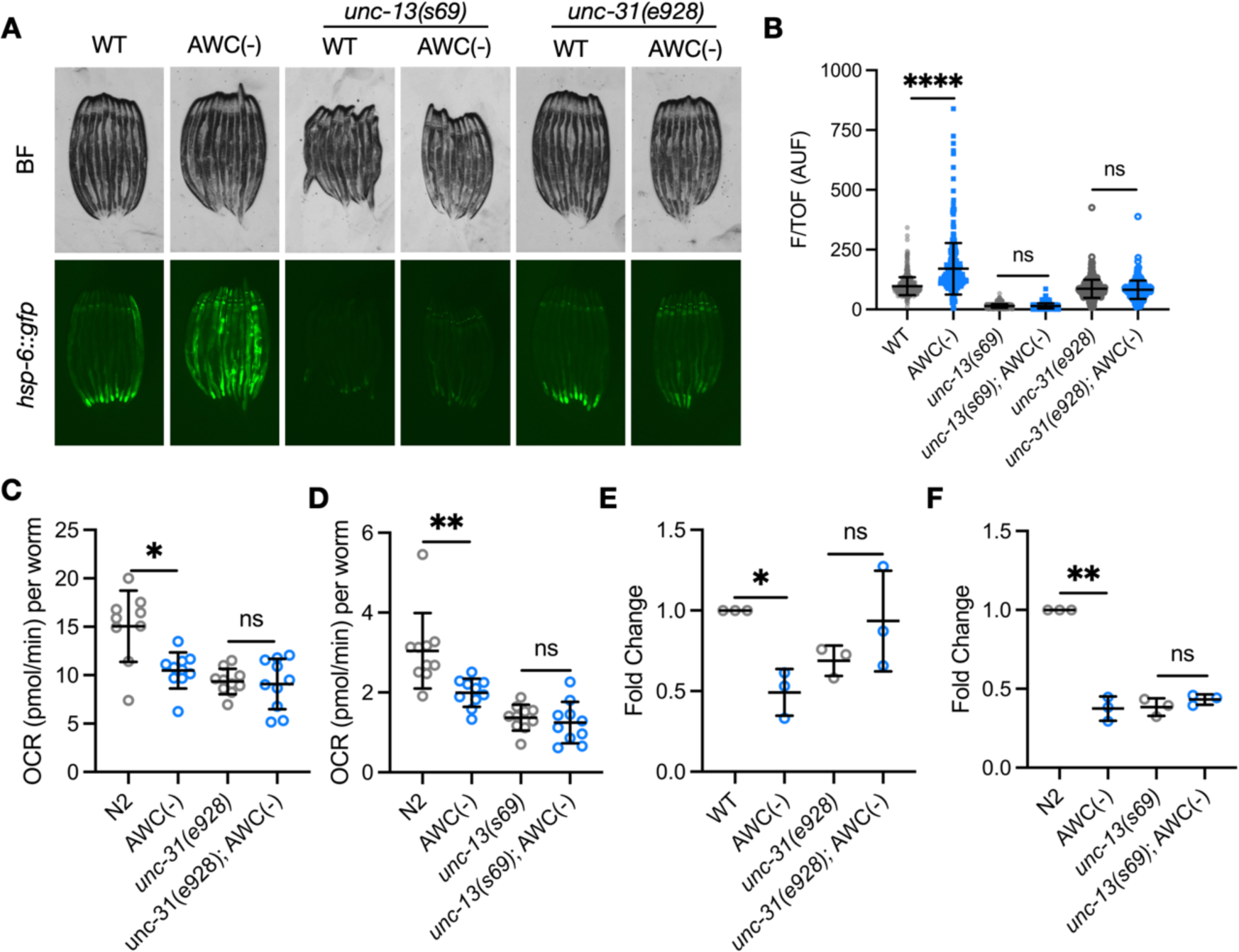
Ablation of AWC olfactory neurons remodels mitochondria through neurotransmission. (A) Representative fluorescent images of *hsp-6::gfp* in N2, AWC(-), *unc-13(s69),* AWC(-); *unc-13(s69), unc-31(e928)* and AWC(-); *unc-31(e928)* animals. (B) Integrated fluorescence intensity measured by bioSorter of *hsp-6::gfp* of strains imaged in panel A. AUF, arbitrary units of fluorescence. Two-tailed unpaired t test with Welch’s correction, ****P<0.0001, P = 0.3813, P = 0.1862. (C) Basal OCR in N2, AWC(-), *unc-31(e928)*, and *unc-31(e928)*; AWC(-) animals. Two-tailed unpaired t test with Welch’s correction, *P<0.05, P = 0.7778. (D) Basal OCR in N2, AWC(-), *unc-13(s69)*, and *unc-13(s69)*; AWC(-) animals. Two-tailed unpaired t test with Welch’s correction, **P<0.01, P = 0.5398. (E) Log2 fold change of the ratio of mtDNA (*mito-1*) to genomic DNA (*ama-1*) of AWC(-), *unc-31(e928)*, and *unc-31(e928)*; AWC(-) normalized to N2 measured by qPCR. N = 3 biological replicates. Two-tailed unpaired t test with Welch’s correction, *P < 0.05, P=0.1178. (F) Log2 fold change of the ratio of mtDNA (*mito-1*) to genomic DNA (*ama-1*) of AWC(-), *unc-13(s69)*, and *unc-13(s69)*; AWC(-) normalized to N2 measured by qPCR. N = 3 biological replicates. Two-tailed unpaired t test with Welch’s correction, **P < 0.01, P = 0.2867.

### Ablation of AWC olfactory neurons remodel mitochondria dependent on serotonin signaling

Having determined that AWC-ablation remodels peripheral mitochondrial dynamics nonautonomously, we investigated what specific molecule(s) carried by either SCVs or DCVs propagate this signal. SCVs carry canonical neurotransmitters and biogenic amines, while DCVs carry biogenic amines and neuropeptides. The biogenic amine serotonin, transmitted in both SCVs and DCVs, has previously been established as a mitokine (*4*, *5*, *20*). Thus, we tested whether loss of *tph-1*, the gene required for the synthesis of serotonin, was required for AWC-mediated UPR^MT^ induction. Indeed, AWC(-); *tph-1(mg280)* mutants completely suppressed UPR^MT^ induction (**Fig. 3, A and B**). Next, we measured OCR and mtDNA in AWC(-); *tph-1(mg280)* mutants and *tph-1(mg280)* mutants, and found that loss of serotonin signaling rescued OCR and mtDNA in AWC(-) animals to wild-type levels (**Fig. 3, C and D**).

**Figure 3.**
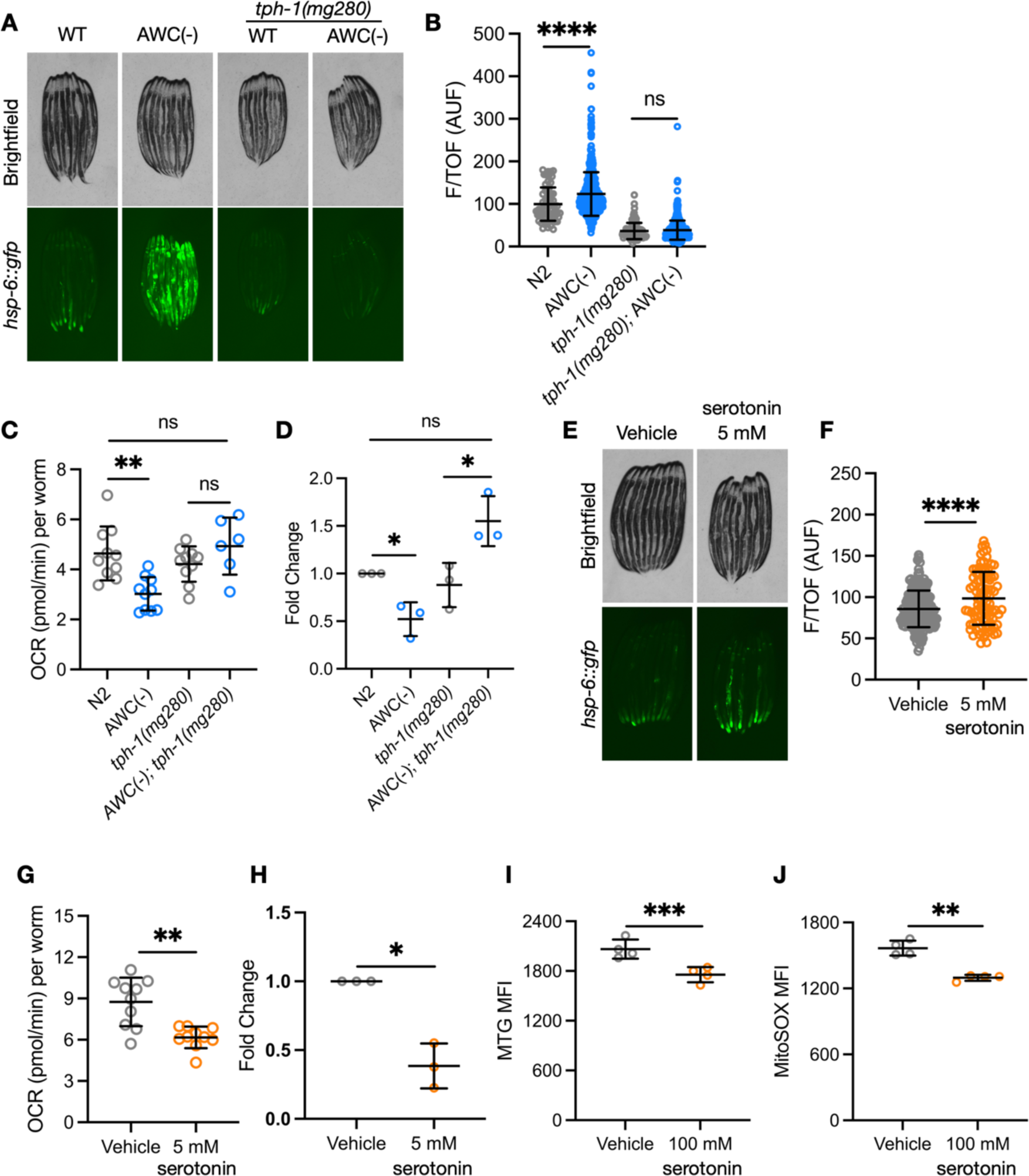
Ablation of AWC olfactory neurons remodels mitochondria dependent on serotonin signaling. (A) Representative fluorescent images of *hsp-6::gfp* in N2, AWC(-), *tph-1(mg280)*, *tph-1(mg280);* AWC(-) animals. (B) Integrated fluorescence intensity measured by bioSorter of *hsp-6::gfp* of strains imaged in panel A. AUF, arbitrary units of fluorescence. Two-tailed unpaired t test with Welch’s correction, ****P<0.0001, P = 0.3670. (C) Basal OCR in N2, AWC(-), *tph-1(mg280)*, and *tph-1(mg280)*; AWC(-) animals. Two-tailed unpaired t test with Welch’s correction, **P<0.01, P = 0.6272, P = 0.2055. (D) Log2 fold change of the ratio of mtDNA (*mito-1*) to genomic DNA (*ama-1*) of AWC(-), *tph-1(mg280)*, and *tphi-1(mg280)*; AWC(-) normalized to N2 measured by qPCR. N = 3 biological replicates. Two-tailed unpaired t test with Welch’s correction, *P < 0.05, P=0.0683. (E) Representative fluorescent images of *hsp-6::gfp* in vehicle and 5 mM serotonin-treated N2 animals. (F) Integrated fluorescence intensity measured by bioSorter of *hsp-6::gfp* of strains imaged in panel E. AUF, arbitrary units of fluorescence. Two-tailed unpaired t test with Welch’s correction, ****P<0.0001. (G) Basal OCR in vehicle and 5 mM serotonin-treated N2 animals. Two-tailed unpaired t test with Welch’s correction, **P<0.01. (H) Log2 fold change of the ratio of mtDNA (*mito-1*) to genomic DNA (*ama-1*) of 5 mM serotonin treated N2 animals normalized to vehicle treated N2 animals measured by qPCR. N = 3 biological replicates. Two-tailed unpaired t test with Welch’s correction, *P < 0.05. (I) Mean fluorescence intensity (MFI) of BJ fibroblasts stained with MitoTracker Green, measured by FACS. Two-tailed unpaired t test with Welch’s correction, ***P < 0.001. (J) Mean fluorescence intensity (MFI) of BJ fibroblasts stained with MitoSOX, measured by FACS. Two-tailed unpaired t test with Welch’s correction, **P < 0.01.

Demonstrating the requirement of serotonin for the changes in mitochondrial dynamics, we examined whether exogenous serotonin treatment was sufficient to recapitulate these phenotypes. First, we treated wild-type UPR^MT^-reporter animals with exogenous serotonin, which was sufficient to induce the UPR^MT^ (**Fig. 3, E and F**). We then treated N2 animals with exogenous serotonin and measured OCR and mtDNA levels. We found that serotonin treatment was also sufficient in reducing OCR and mtDNA (**Fig. 3, G and H**). Together, these data indicate that AWC-ablation remodels peripheral mitochondrial dynamics through disinhibited serotonergic signaling.

To more directly test whether serotonin signaling regulates mitochondrial homeostasis in recipient cells, we supplemented serotonin to cultured human BJ fibroblasts, a karyotypically stable cell line that naturally expresses multiple serotonin receptors (*21*). Consistent with what we observed in *C. elegans* models, serotonin-treated cells had reduced MitoTracker Green staining, indicative of reduced mitochondrial mass (**Fig. 3I**). Moreover, serotonin treatment also reduced MitoSOX Red staining, indicating that the generation of mitochondrial reactive oxygen species (ROS) and probably overall mitochondrial electron transport chain activities are declined (**Fig. 3J**). These data demonstrate that serotonin may have a conserved role in regulating mitochondrial function across species.

### Ablation of AWC remodels mitochondrial dynamics dependent on PDR-1

Having observed that AWC(-) animals have reduced mtDNA, we tested if the induced UPR^MT^, reduced OCR, and reduced mtDNA phenotypes were driven by mitophagy. To test this, we measured whether these mitochondrial phenotypes depend on *pdr-1*/*Parkin*. First, we compared mtDNA in AWC(-); *pdr-1(gk448)* mutants to AWC(-) mutants, and observed a rescue in mtDNA levels (**Fig. 4A**). Additionally, we observed a rescue in OCR in AWC(-); *pdr-1(gk448)* compared to AWC(-) mutants (**Fig. 4B**). Next, we crossed *pdr-1(gk448)* to AWC(-); *hsp-6::gfp* mutants, and observed suppression of UPR^MT^ induction (**Fig. 4, C and D**). We were surprised by this result, as it suggests that induction of the UPR^MT^ occurs downstream of PDR-1-dependent activity. To determine whether *atfs-1*, the transcriptional regulator of the UPR^MT^, drives the OCR and mtDNA phenotypes, we measured OCR and mtDNA levels in AWC(-); *atfs-1(gk3094)* mutants compared to AWC(-) mutants. Interestingly, loss of *atfs-1* further reduced OCR in AWC(-) and did not rescue mtDNA levels in AWC(-) to WT levels (**Fig. 4, E and F**). We interpreted this result to suggest that AWC-ablation reduces mtDNA and OCR in a PDR-1-dependent manner, which may subsequently drive ATFS-1 localization to the nucleus to induce the UPR^MT^.

**Figure 4.**
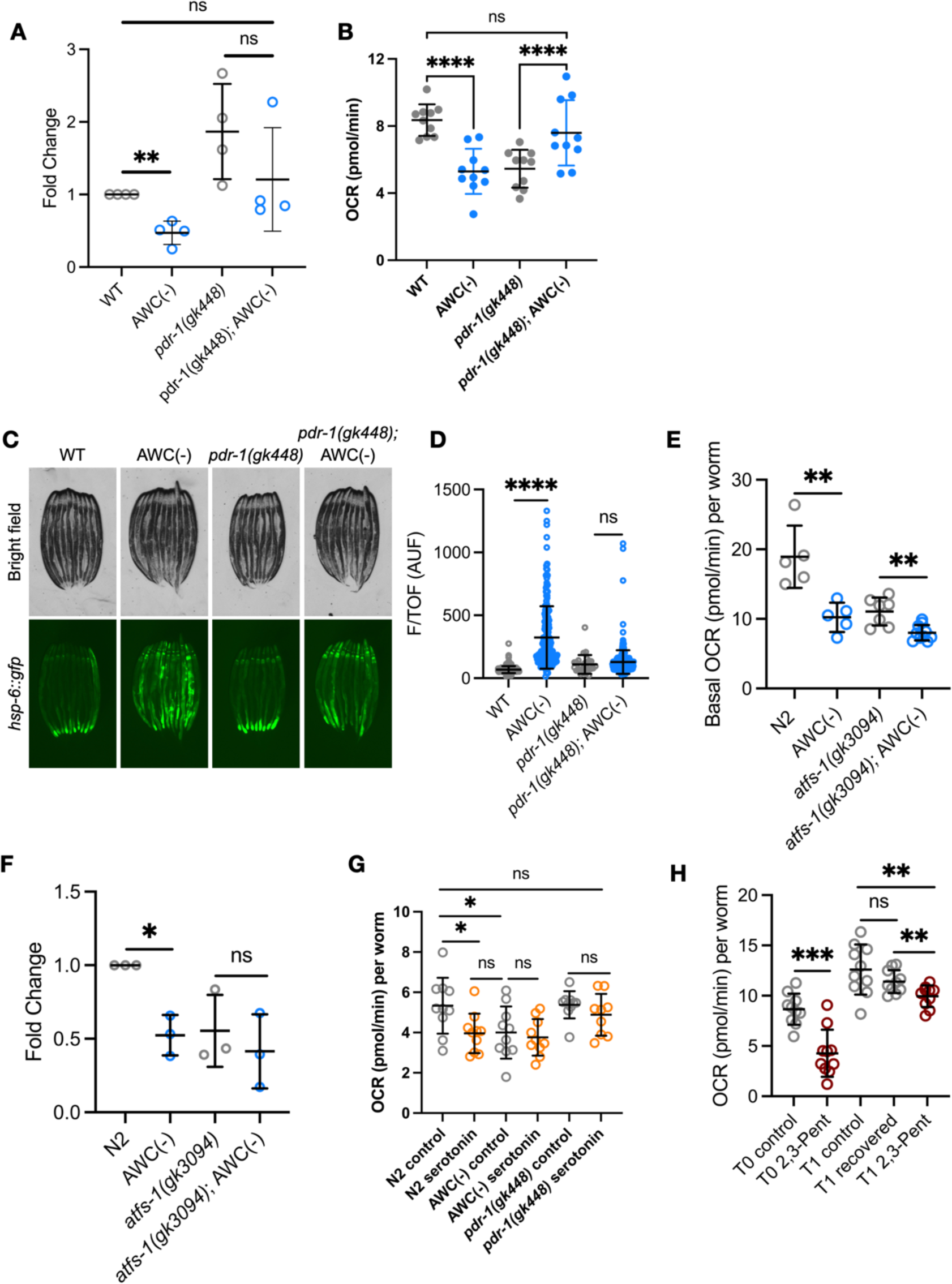
Ablation of AWC remodels mitochondrial dynamics dependent on PDR-1. (A) Log2 fold change of the ratio of mtDNA (*mito-1*) to genomic DNA (*ama-1*) of N2, AWC(-), *pdr-1(gk448)* and *pdr-1(gk448)*; AWC(-) strains normalized to N2 animals, measured by qPCR. N = 4 biological replicates. Two-tailed unpaired t test with Welch’s correction, **P < 0.01. P = 0.2227, P = 0.0776. (B) Basal OCR in N2, AWC(-), *pdr-1(gk448)* and *pdr-1(gk448)*; AWC(-) animals. Two-tailed unpaired t test with Welch’s correction, ****P<0.001, P = 0.2877. (C) Representative fluorescent images of *hsp-6::gfp* in N2, AWC(-), *pdr-1(gk448)* and *pdr-1(gk448)*; AWC(-) animals. (D) Integrated fluorescence intensity of *hsp-6::gfp* of strains imaged in panel C. AUF, arbitrary units of fluorescence. Two-tailed unpaired t test with Welch’s correction, ****P<0.0001, P = 0.2932. (E) Basal OCR in N2, AWC(-), *atfs-1(gk3094)*, and *atfs-1(gk3094)*; AWC(-) animals. Two-tailed unpaired t test with Welch’s correction, **P<0.01. (F) Log2 fold change of the ratio of mtDNA (*mito-1*) to genomic DNA (*ama-1*) of N2, AWC(-), *atfs-1(gk3094)*, and *atfs-1(gk3094)*; AWC(-) strains normalized to N2 animals, measured by qPCR. N = 3 biological replicates. Two-tailed unpaired t test with Welch’s correction, *P < 0.05. P = 0.5298. (G) Basal OCR in N2, AWC(-), and *pdr-1(gk448)* animals treated with vehicle or serotonin. Two-tailed unpaired t test with Welch’s correction, *P<0.05. (H) Basal OCR in control and 2,3-Pent treatment at L4 (T0). Basal OCR in control, 24h 2,3-Pent recovery, and chronic 2,3-Pent exposure (T1). Two-tailed unpaired t test with Welch’s correction, ***P<0.001, **P<0.01, P = 0.1910.

To understand if serotonin signaling reduces OCR and mtDNA levels dependent on *pdr-1*, we treated N2, AWC(-), and *pdr-1(gk448)* mutants with exogenous serotonin. We observed that serotonin treatment failed to further reduce OCR in AWC(-) mutants, and that serotonin failed to reduce OCR in *pdr-1(gk448)* mutants (**Fig. 4G**). These data suggest that PDR-1 functions downstream of serotonin activity to reduce OCR and mtDNA.

Finally, we asked whether silencing of AWC permanently or transiently influences mitochondrial function. To test this, we exposed N2 animals to 2,3-Pent from hatch until the L4 stage (T0), then allowed them to recover for 24h in the absence of odorant (T1). As previously shown, 2,3-Pent exposure through the L4 stage was sufficient to reduce OCR (**Fig. 4H**). We found that a 24h recovery after L4 rescued OCR to wild-type levels compared to N2 animals that were chronically 2,3-Pent exposed (**Fig. 4H**). This suggests that 2,3-Pent exposure transiently reduces OXPHOS, and re-activation of AWC by odorant removal promotes OXPHOS recovery, perhaps through compensatory mitochondrial biogenesis.

### Ablation of AWC confers pathogen resistance dependent on ATFS-1 and PDR-1

Ablation of AWC increases resistance to several pathogens, including the human pathogen *Pseudomonas aeruginosa* (strain PA14), but the mechanism by which AWC ablation confers pathogen resistance is unknown (*12*). During infection with PA14, *C. elegans* activates *atfs-1* to resolve pathogen-driven mitochondrial damage and to induce innate immune response genes (*11*). Further, loss of *atfs-1* and diminished UPR^MT^ induction results in sensitivity to PA14 infection (*11*). Thus, we asked whether the observed UPR^MT^ phenotype in AWC(-) might contribute to this resistance. To test this, we exposed AWC(-) and AWC(-); *atfs-1* to PA14, and observed that AWC(-) mutants were resistant to PA14 partially depending on *atfs-1* (**Fig. 5A**). Mitophagy has been identified as a pro-survival strategy during infection by *P. aeruginosa*, and mutants lacking mitophagy machinery are more sensitive to *P. aeruginosa* infection (*22*). In an effort to proliferate, *P. aeruginosa* chelates rate-limiting iron from host mitochondria, and in response, *C. elegans* performs mitophagy to limit *P. aeruginosa* virulence (*22*). Because UPR^MT^ induction in AWC(-) mutants was PDR-1-dependent, we hypothesized that induction of mitophagy downstream of AWC-ablation may contribute to *P. aeruginosa* resistance. We performed a PA14 survival assay on N2, AWC(-), *pdr-1(gk448)*, and *pdr-1(gk448)*; AWC(-) strains, and found that the survival benefit of AWC(-) animals fully depends on *pdr-1* (**Fig. 5B)**.

**Figure 5:**
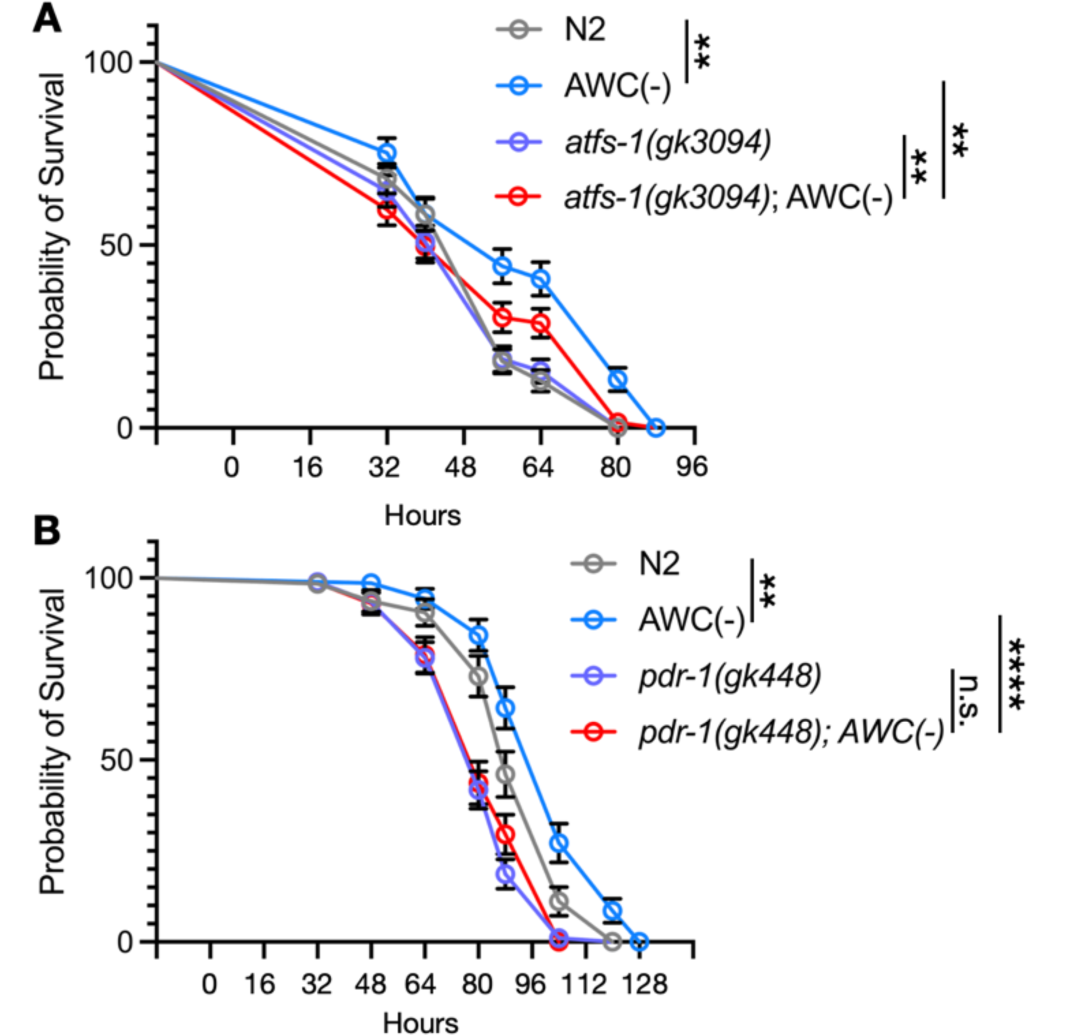
Ablation of AWC confers pathogen resistance dependent on ATFS-1 and PDR-1. (A) Survival of N2, AWC(-), *atfs-1(gk3094)*, and *atfs-1(gk3094)*; AWC(-) on PA14. Log-rank Mantel Cox test, **P<0.01. (B) Survival of N2, AWC(-), *pdr-1(gk448)*, and *pdr-1(gk448)*; AWC(-) on PA14. Log-rank Mantel Cox test, ****P<0.001, **P<0.01, P = 0.3410.

## DISCUSSION

We have identified a unique role for AWC olfactory neurons in regulating peripheral mitochondrial dynamics. AWC(-) animals exhibit induction of the UPR^MT^, reduction in OXPHOS, and a depletion of mtDNA dependent on serotonergic signaling (Figure 1). These data fit with previous studies that established serotonin as a mitokine, where serotonergic signaling was required for UPR^MT^ activation downstream of neuronal mitochondrial dysfunction (*4*, *5*, *20*). Here, however, we also show that exogenous serotonin treatment is sufficient to drive the UPR^MT^, reduce OXPHOS, and reduce mtDNA, phenocopying AWC ablation (Figure 3). This suggests that AWC ablation may drive these mitochondrial phenotypes through disinhibited serotonin signaling. We also demonstrate that serotonin may play a conserved role in mitochondrial regulation in mammalian cells, whereby serotonin treatment reduces mitochondrial mass and activity in mammalian fibroblasts (Figure 3). In the broader context, there is evidence that neuropeptidergic afferents to the mesentary can drive the release of serotonin from mast cells, which store large quantities of the neurotransmitter (*26*). In the intestine, serotonin is an important signaling molecule that drives immune cell activation, the release of pro-inflammatory cytokines, increases cytotoxicity, and is important for maintaining microbiome homeostasis (*27*). Therefore, we hypothesize if a similar olfactory-to-intestine signaling axis is conserved in mammals, whereby olfactory activation may increase intestinal serotonin activity to increase immune activation.

Further, we show that the depletion of mtDNA and OXPHOS phenotypes observed in AWC(-) are dependent on mitophagy machinery *pdr-1/Parkin* (Figure 4). We determined that serotonin treatment reduces OXPHOS in a *pdr-1*-dependent fashion, suggesting that serotonin may drive enhanced mitophagy, and that AWC ablation may drive mitophagy through serotonergic signaling. Induction of the UPR^MT^ in AWC(-) animals also depends on *pdr-1*, and we find that 2,3-Pentanedione can reduce OXPHOS and mtDNA levels before the activation of the UPR^MT^. Previous studies have shown that mitophagy occurs in response to mitochondrial dysfunction and UPR^MT^ induction (*25*), but here we show that mitophagy may also be able to precede *atfs-1* activation.

Previous studies have sought to understand whether specific neurons, particularly sensory neurons, were able to coordinate peripheral UPR^MT^ and mitochondrial function. Overexpression of the mitokine Wnt/EGL-20 in ADL gustatory neurons was sufficient to drive peripheral UPR^MT^ induction and pathogen survival depending on *atfs-1* (*23*). Here, however, we show that the nervous system can modulate peripheral mitochondrial dynamics in response to a naturalistic cue, in addition to genetic activation of neuronal mitochondrial dysfunction. Specifically, the naturalistic AWC-silencing odorant 2,3-Pentanedione is sufficient to reduce OXPHOS and induce the UPR^MT^, while also showing a trend in depleting mtDNA. 2,3-Pentanedione is a metabolite in the biosynthetic process of the universal autoinducer-2 (AI-2), an important bacterial quorum sensing molecule (*24*). We speculate that the olfactory nervous system may have evolved a unique role in relaying odorant cues, like 2,3-Pentanedione, to prepare the periphery for potential homeostatic perturbations, such as metabolic stress or pathogenic infection.

Finally, we validate that AWC(-) animals are resistant to infection with the pathogenic bacteria *P. aeruginosa* (*12*) and that this resistance is partially dependent on *atfs-1*, and fully dependent on *pdr-1*. *P. aeruginosa* exploits *C. elegans* mitochondria as an iron source through production of the iron chelator pyoverdine, and loss of the ability to perform mitophagy sensitized *C. elegans* to pathogenic infection (*22*). This study speculated that *C elegans* evolved to perform mitophagy to limit *P. aeruginosa* virulence by limiting its iron source. With these data, we hypothesize that olfactory neurons may couple bacterial odorant cues to mitophagy as an anticipatory strategy to evade pathogenic insult.

## MATERIALS AND METHODS

### *C elegans* strains and details

Nematodes were maintained at 15°C or 20°C on standard nematode growth medium (NGM) agar plates seeded with *Escherichia coli* strain OP50. All experiments were conducted on bleach synchronized populations, at 20°C on OP50 plates. Synchronization was achieved by washing animals fed with OP50 with M9 solution (22 mM KH2PO4 monobasic, 42.3 mM Na2HPO4, 85.6 mM NaCl, and 1 mM MgSO4), bleached using a solution of 1.8% sodium hypochlorite and 0.375 M KOH diluted in DDW, for 4-5 minutes. Intact eggs were then washed 3× with M9 solution, and intact eggs were verified under the microscope after seeding.

### Microscopy and quantification of *hsp-6::GFP* reporter

Imaging of strains in the *hsp-6::GFP* background was performed as previously described (Bar-Ziv et al., 2020). Briefly, strains were grown from egg to day 1 of adulthood (D1), unless otherwise specified, at 20°C and picked in bright field to avoid biasing reporter intensity. Animals were paralyzed with 10µL of 100mM sodium azide and oriented from head to tail on unseeded NGM plates. Images were captured on a Leica M250FA stereoscope with a Hamamatsu ORCA-ER camera with LAS-X-software. Bright field and fluorescent images were taken with experiment-matched exposure time and laser intensity with a 1X objective and 4X magnification. Each microscopy experiment was sampled from a population that was subsequently quantified using a large-particle flow cytometer (Union Biometrica bioSorter) as previously described (Bar-Ziv, 2020). Data for time of flight (TOF, a measure of length), extinction (width) and integrated GFP fluorescence was collected. Data were gated by TOF and extinction to exclude eggs or any larvae present. Data are presented as integrated fluorescence divided by TOF, to normalize for varying lengths (F/TOF) in arbitrary units (AUF). All experiments contained N > 250 worms.

### Oxygen consumption rate (OCR) measurement

OCR was measured using a Seahorse XeF96 (Agilent) instrument using the extracellular flux assay kit (Agilent). 20-40 L4 animals were plated in 200µL of M9 per well, with 5-10 wells per strain. Basal oxygen consumption rate was read using the following program: 5 cycles of 2 minutes mix, 30 seconds wait, 2 minutes measure. For maximal oxygen consumption, 22µL of 100 mM carbonyl cyanide 4-(trifluoromethoxy)phenylhydrazone (FCCP; Sigma Aldrich) was injected into each well, and OCR was measured using the following program: 9 cycles of 2 minutes mix, 30 seconds wait, 2 minutes measure. All assays ended with sodium azide (NaN3) treatment to measure non-mitochondrial respiration, and to paralyze the worms for quantification. 22µL of 400 mM NaN3 was injected into each well and was measured using the following program: 4 cycles of 2 minutes mix, 30 seconds wait, and 2 minutes measure. Following the assay, paralyzed worms were quantified per well. In OCR figures, each data point represents the average oxygen consumption rate in pmol per minute per the quantity of worms in each well.

### Mitochondrial DNA (mtDNA) quantification

mtDNA was quantified by quantitative PCR (qPCR) as previously described (Shpilka et al., 2020). Briefly, 20-30 L4 animals were collected in 20 µL of lysis buffer (50 mM KCl, 10 mM Tris-HCl pH 8.3, 2.5 mM MgCl2, 0.45% NP-40, 0.45% Tween 20, 0.01% gelatin, with freshly added 200 µg/ml proteinase K) and flash frozen at −80°C for at least 20 minutes prior to lysis. Lysis was performed at 65°C for 80 minutes. Each reaction was done in triplicate using 1µL of mtDNA lysate, with SYBR Select Master Mix (Thermo Fisher Scientific). mtDNA-specific primers used: *mito-1*: F’ *gtttatgctgctgtagcgtg*, R’ *ctgttaaagcaagtggacgag*; *nd-1*: F’ *agcgtcatttattgggaagaagac*, R’ *aagcttgtgctaatcccataaatgt*. Genomic-DNA-specific primers used: *ama-1*: F’ *tggaactctggagtcacacc*, R’ *catcctccttcattgaacgg*; *cox-4*: F’ *gccgactggaagaacttgtc*, R’ *gcggagatcaccttccagta*. Data presented are ratios of mtDNA to genomic DNA (*mito-1*:*ama-1* or *nd-1*:*cox-4*) normalized to N2 values, with N = 3 biological replicates.

### Odorant exposure

Bleach synchronized eggs were seeded onto OP50 plates. For odorant conditions, 1µL of undiluted odorant was applied to the lid of the plate and inverted. Odorant-exposed plates were placed in a 3×5 in. air-tight box, in a fume hood at RT. Control condition worms were placed in an identical box without odors in the same fume hood. Odors were refreshed every 24h. Odorants used were 2,3-Pentanedione (2,3-Pent; Sigma Aldrich) and 2-Butanone (2-Bt; Sigma Aldrich).

### Pharmacological treatments in *C elegans*

To inhibit germline proliferation, L4 populations were washed off plates in M9 and seeded onto pre-treated (+)-5-Fluorodeoxyuridine (FUDR; Spectrum Chemical) OP50 plates. Plates were pre-treated with 100µL of FUDR [10mg/mL] and allowed to dry before L4 worms were plated. For assays using the AWC-specific histamine-gated chloride channel construct (*ceh-36p::HisCl*), OP50 plates were treated with 500µL of 200 mM histamine dihydrochloride (HA; Sigma Aldrich) dissolved in water, for a final plate concentration of 10 mM. L4 *ceh-36p::HisCl* worms were transferred to dried HA or vehicle-treated plates and assayed at D1.

For exogenous serotonin treatment, OP50 plates were treated with 330µL of 150 mM serotonin hydrochloride (Sigma Aldrich), dissolved in water, for a final plate concentration of 5 mM. Synchronized eggs were plated onto dried vehicle-treated or serotonin-treated plates. qPCR and OCR for serotonin-treated worms were performed at L4, while *hsp-6::gfp* imaging was performed at D1 of adulthood.

### Cell Culture

Cells were grown in DMEM media (11995, Thermo Fisher) supplemented with 2 mM GlutaMAX (35050, Thermo Fisher), 10% FBS (VWR), Non-Essential Amino Acids (100 X, 11140, Thermo Fisher), and Penicillin-Streptomycin (100 X, 15070, Thermo Fisher) in 5% CO2 at 37 °C. 293T cells were obtained from UC Berkeley Cell Culture Facility and hTERT-immortalized human foreskin BJ fibroblasts were obtained from ATCC (CRL-4001). The identity of cell lines were confirmed through their STR profling by UC Berkeley DNA Sequencing Facility. Mycoplasma negative status confirmed by PCR Detection Kit.

### Flow Cytometry

Cells were plated onto culture plates (Corning) and allowed to grow overnight. Drugs were directly added to media. Cells (should be around 80% confluency) were harvested after 24 hours of the drug treatment. To perform flow cytometry staining, cells were trypsinized and stained with 150 nM MitoTracker Green (M7514, Thermo Fisher) or 5 µM MitoSOX Red (M36008, Thermo Fisher) in culture media at 37 °C for 30 min. After staining, cells were washed once with cold culture media and resuspended in cold FACS buffer (PBS with 0.1% BSA and 2 mM EDTA). Samples were then analyzed with a four-laser Attune NxT flow cytometer (Thermo Fisher). Acquired data were analyzed using FlowJo 10.

### *P. aeruginosa* survival assay

*P. aeruginosa* strain PA14 was cultured at 37°C in LB overnight before spotting 20µL of inoculum onto slow killing plates. Inoculum was spread to cover >75% of the plate. Seeded plates were cultured at 37°C overnight. PA14 plates were allowed to come to room temperature before treatment with FUDR as previously described. FUDR was used to prevent pathogen-induced intestinal egg hatching. Synchronized L4 worms were transferred to PA14 plates, with 10-15 worms on 8 plates. Lethality was determined by the absence of movement after a nose and tail prod. Lethality was measured each day, with more frequent measurements as survival decreased.

### 2,3-Pentanedione (2,3-Pent) odorant recovery assay

Synchronized eggs were plated on OP50 and treated with 2,3-Pent or no odorant as described. Baseline OCR measurements were performed at L4 stage. 2,3-Pent-treated L4 animals were washed off plates using M9, and washed 3x with M9 before re-plating on a fresh OP50 plate. 2,3-Pent-exposed animals were allowed to recover without odor for 24h. OCR was performed on 2,3-Pent-recovered animals, no odor-exposed animals, and continuously 2,3-Pent-exposed animals at D1 of adulthood.

### Statistical analyses

Statistical analysis was performed using GraphPad Prism 9.2.0. Individual analyses are as described in figure legends. Lifespans were analyzed using a log-rank Mantel Cox test. Two condition comparisons were analyzed using two-tailed unpaired Welch’s t tests.

---

**Supplemental Figure 1.**
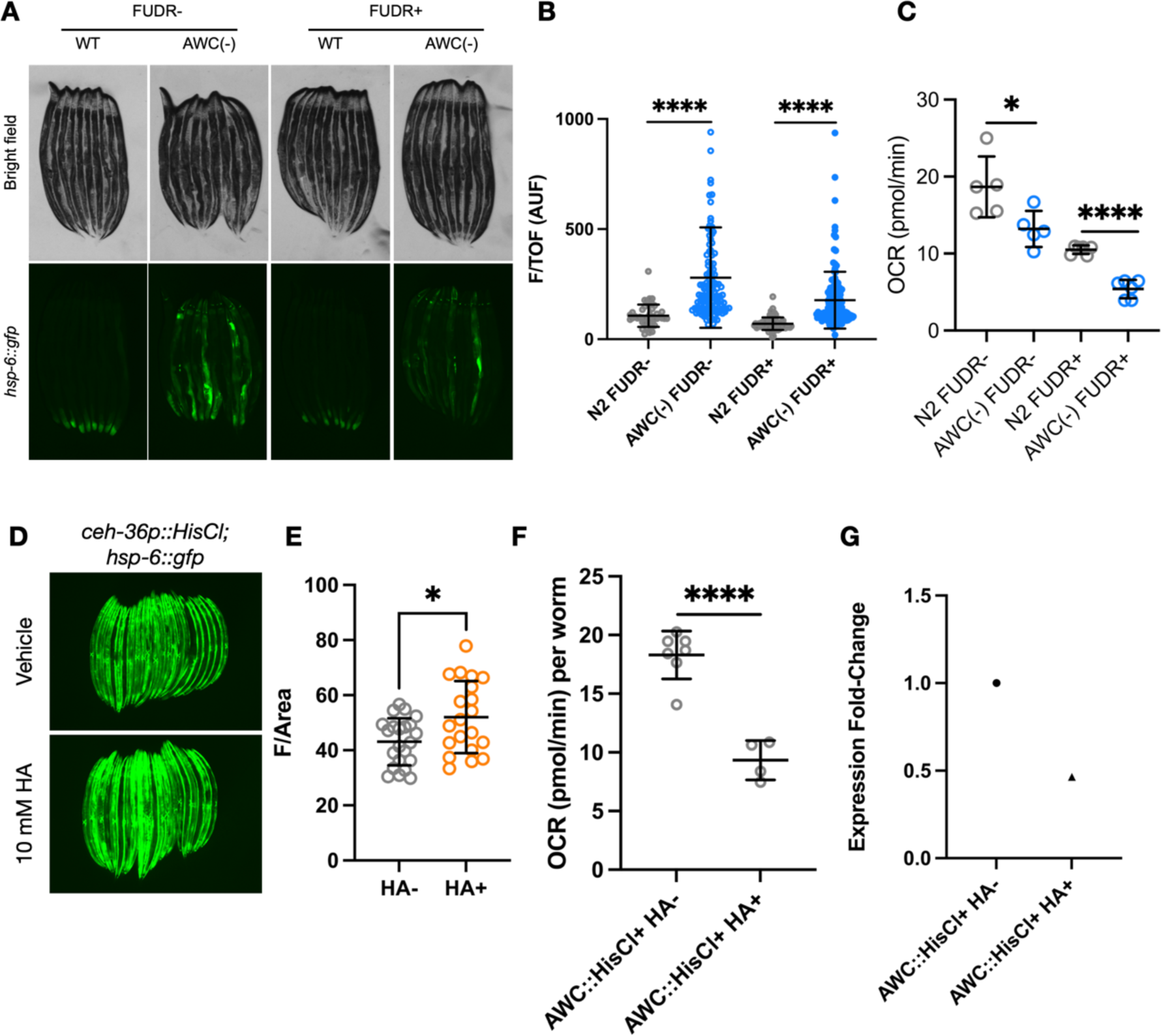
(A) Representative fluorescent images of *hsp-6::gfp* in N2 and AWC(-) without and with FUDR. (B) Integrated fluorescence intensity of *hsp-6::gfp* of strains imaged in panel A. AUF, arbitrary units of fluorescence. Two-tailed unpaired t test with Welch’s correction, ****P<0.0001. (C) Basal OCR in N2 and AWC(-) without and with FUDR. Two-tailed unpaired t test with Welch’s correction, *P<0.01, ****P<0.0001. (D) Representative fluorescent images of *hsp-6::gfp* in *ceh-36p::HisCl* animals treated without or with 10 mM histamine (HA) (E) Fluorescence intensity of intestinal *hsp-6::gfp* expression measured by ImageJ between HA- and HA+ *ceh-36p::HisCl* animals. *P<0.05. (F) Basal OCR in in *ceh-36p::HisCl* with or without 10 mM HA treatment. Two-tailed unpaired t test with Welch’s correction, ****P<0.0001. (F) Log2 fold change of the ratio of mtDNA (mito-1) to genomic DNA (ama-1) of *ceh-36p::HisCl* with 10 mM HA treatment normalized to no HA treatment, measured by qPCR. N = 1 biological replicates.

